# Kinetic modeling predicts a role for ribosome collisions at elongation stall sites in bacteria

**DOI:** 10.1101/087668

**Authors:** Michael Ferrin, Arvind R. Subramaniam

## Abstract

Ribosomes can stall during translation elongation in bacteria and eukaryotes. To identify mechanisms by which ribosome stalling affects expression of the encoded protein, we develop an inverse approach that combines computational modeling with systematic perturbations of translation initiation rate, the number of stall sites, and the distance between stall sites on a reporter mRNA. By applying this approach to ribosome stalls caused by amino acid starvation in the bacterium *Escherichia coli*, we find that our measurements are quantitatively inconsistent with two widely used kinetic models for stalled ribosomes: ribosome traffic jams that block initiation, and abortive (premature) termination of stalled ribosomes. To account for this discrepancy, we consider a model in which collision from a trailing ribosome causes abortive termination of the stalled ribosome. This collision-stimulated abortive termination model provides a better fit to measured protein synthesis rates from our reporter library, and is consistent with observed ribosome densities near stall sites. Analysis of this model further predicts that ribosome collisions can selectively stimulate abortive termination of stalled ribosomes without fine-tuning of kinetic rate parameters. Thus ribosome collisions may serve as a robust timer for translational quality control pathways to recognize stalled ribosomes.

## Introduction

Ribosomes move at an average speed of 3–20 codons per second during translation elongation *in vivo* (1–3). Since this rate is higher than the typical initiation rate of ribosomes on mRNAs [less than 1s^-1^ (3, 4)], elongation is often assumed to not affect the expression level of most proteins. Nevertheless, elongation rate of ribosomes can decrease significantly at specific locations on an mRNA due to low abundance of aminoacyl-tRNAs, inhibitory codon pairs or amino acid pairs, nascent peptides interacting strongly with the ribosome exit tunnel, and presence of RNA-binding proteins (5). Ribosome profiling — the deep sequencing of ribosome-protected mRNA fragments — has enabled the identification of several factors that induce slowing or stalling of ribosomes during elongation (6, 7). An important question emerging from these studies is the extent to which ribosome stalling affects the expression of the encoded protein, since initiation might still be the slowest step during translation.

Several mechanistic models have been proposed to explain how ribosome stalling during elongation might affect the expression of the encoded protein. In the widely used traffic jam model (8), the duration of ribosome stalling is sufficiently long to induce a queue of trailing ribosomes extending to the start codon, thus decreasing the translation initiation rate. Evidence supporting this model has been found in the context of EF-P dependent polyproline stalls in *E. coli* (9, 10), and rare-codon induced pausing in *E. coli* and yeast (11, 12). In an alternate abortive termination model, ribosome stalling causes premature termination without synthesis of the full-length protein. This model is thought to underlie the action of various ribosome rescue factors in *E. coli* and yeast (13, 14). Finally, ribosome stalling can also affect protein expression indirectly by altering mRNA stability (15, 16), co-translational protein folding (17), or stress-response signaling (18).

Despite the experimental evidence supporting the above models, predicting the effect of ribosome stalling on protein levels has been challenging because of uncertainty in our knowledge of *in vivo* kinetic parameters such as the duration of ribosome stalling and the rate of abortive termination. Further, while we have a detailed understanding of the kinetic steps and structural changes that occur during the normal elongation cycle of the ribosome (19–21), the ‘off-pathway’ events that occur at stalled ribosomes have been elucidated in only a few specific cases (22–24). Thus, development of complementary approaches, which can quantitatively constrain the *in vivo* kinetics of stalled ribosomes without precise knowledge of rate parameters, will be useful for bridging the gap between the growing list of ribosome stall sequences (7, 25, 26) and their effect on protein expression.

Here, we investigated the effect of ribosome stalling on protein expression using amino acid starvation in *E. coli* as an experimental system. In this system, we had previously found that both ribosome traffic jams and abortive termination occur at a subset of codons cognate to the limiting amino acid (27). Motivated by these observations, here we computationally modeled ribosome traffic jams and abortive termination with the goal of predicting their effect on protein expression. Even without precise knowledge of *in vivo* kinetic parameters, we found that these two processes give qualitatively different trends in protein expression when the initiation rate, the number of stall sites and the distance between stall sites are systematically varied. Surprisingly, experimental measurements support a model in which traffic jams and abortive termination do not occur independent of one another; rather, collisions by trailing ribosomes stimulate abortive termination of the stalled ribosome. Our integrated approach, developed here in the specific context of amino acid starvation in *E. coli*, should be generally applicable to investigate ribosome stalls in both bacteria and eukaryotes.

## Results

### Effect of ribosome stalling on measured protein level, mRNA level and polysome occupancy

During starvation for single amino acids in *E. coli*, certain codons that are cognate to the limiting amino acid decrease protein expression, while the same codons have little or no effect during nutrient-rich growth (28). For example, synonymously mutating seven CTG leucine codons in the yellow fluorescent protein gene (*yfp*) to CTA, CTC or CTT reduced the synthesis rate of YFP 10–100 fold during leucine starvation (28). Genome-wide ribosome profiling showed that ribosomes stall at CTA, CTC and CTT codons during leucine starvation, which leads to a traffic jam of trailing ribosomes and abortive termination of translation (27). These observations led us to ask whether ribosome traffic jams and abortive termination can quantitatively account for the decrease in protein synthesis rate (number of full proteins produced per unit time) caused by ribosome stalling during leucine starvation in *E. coli*.

To measure the effect of ribosome stalling on protein synthesis during leucine starvation, we constructed fluorescent reporter genes which have a stall-inducing CTA codon at one or two different locations along *yfp* (Fig. 1A, blue bars). We induced these reporter variants from very low copy vectors (SC*101 *ori*, 3-4 copies per cell) either during leucine starvation or during leucine-rich growth. While YFP expression was similar across all *yfp* variants during leucine-rich growth, a single CTA codon at two different locations reduced YFP expression during leucine starvation by 3–4 fold relative to a control *yfp* without CTA codons (Fig. 1B). Introducing both CTA codons reduced YFP expression by ~6-fold, and a stretch of 7 CTA codons reduced YFP expression close to background level as observed in our earlier work (28). Thus, YFP expression can serve as a quantitative readout of the effect of ribosome stalling on protein synthesis.

**Figure 1:**
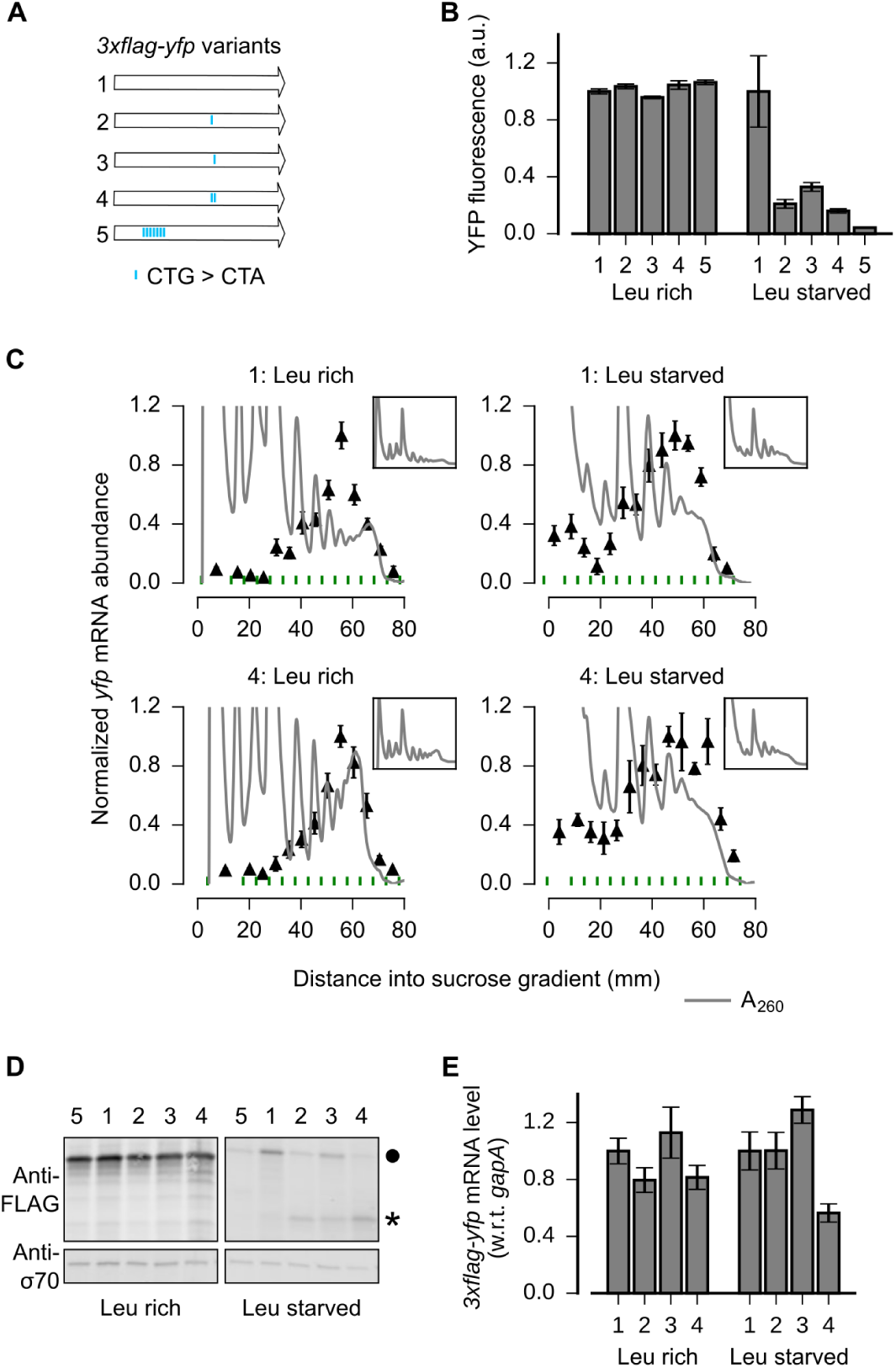
Effect of ribosome stalling on measured protein level, mRNA level and polysome occupancy. **A**. Schematic of ribosome stalling reporters. Blue vertical lines along the 260 codon *3xflag-yfp* reporters indicate location of CTA Leu codons that cause ribosome stalling during Leu starvation in *E. coli*. For experiments in B–E, reporters were induced either in Leu rich growth medium for 20 min or Leu starvation medium for 60 min. **B**. YFP fluorescence normalized by that of construct 1 in each condition. Error bars represent standard error over triplicate cultures. **C**. ▲ indicate *yfp* mRNA levels in polysomes fractionated through a sucrose gradient, and measured by quantitative RT-PCR spanning the CTA codons. Absorbance at 260nm is shown in grey with magnified Y axis to highlight polysome profiles. *Inset*: Polysome profiles showing full monosome peak. Individual polysome fractions are separated by green lines above X axis. An *in vitro* transcribed luciferase mRNA was spiked in for normalizing the mRNA levels by each fraction’s volume. Error bars represent standard error of qPCR over triplicates. **D**. *Top panel*: Western blot against the 3xFLAG epitope at the N-terminus of reporters. ● indicates the size of the full length 3xFLAG-YFP product. ★ indicates the size of truncated product expected from abortive termination of ribosomes at CTA codons in constructs 2, 3 and 4. *Bottom panel*: Western blot against the RNA polymerase σ_70_ subunit shown as a loading control. **E**. *yfp* mRNA levels normalized by that of construct 1 in each condition. *gapA* endogenous mRNA used for internal normalization. Error bars represent standard error of qPCR over triplicates.

We then sought biochemical evidence supporting a role for either ribosome traffic jams or abortive termination in the reduction of YFP expression caused by stall-inducing CTA codons. We reasoned that ribosome traffic jams that reduce protein expression by blocking initiation should increase the number of ribosomes on an mRNA when the stall site is far from the initiation region. However, polysome fractionation of leucine-starved *E. coli* did not indicate an unambiguous shift of the *yfp* mRNA to higher polysome fractions when two stall-inducing CTA codons were introduced >400nt from the start codon (Fig. 1C, top vs bottom panels). This observation agrees with ribosome density measurements that detected a traffic jam of at most 1–2 ribosomes behind the stalled ribosome (27).

We detected truncated YFP fragments consistent with abortive termination at stall-inducing CTA codons during leucine starvation (Fig. 1D). Previous studies suggested that abortive termination of stalled ribosomes requires cleavage of mRNA near the stall site as an obligatory step (29, 30, 13). Therefore, we tested whether changes in mRNA levels could account for the 3-4 fold decrease in YFP expression caused by single CTA codons during leucine starvation (Fig. 1B). However, we found that *yfp* mRNA levels, as measured by quantitative RT-PCR spanning the single CTA codons, did not decrease significantly during leucine starvation (Fig. 1E). Similarly, introducing two CTA codons resulted in <2-fold decrease in *yfp* mRNA levels despite ~6-fold decrease in YFP expression (Fig. 1E vs 1B). These observations are consistent with earlier measurements using ribosome profiling and Northern blotting that did not find evidence for significant mRNA cleavage or decay upon ribosome stalling at CTA codons during leucine starvation (27).

### Computational modeling of ribosome kinetics at stall sites

Since the above reporter-based experiments were qualitative and could miss subtle effects, we formulated an alternate approach using computational modeling to quantitatively test the role of ribosome traffic jams and abortive termination at stall sites. To this end, we defined a minimal set of five kinetic states at ribosome stall sites and the rate constants for transition between these kinetic states (Fig. 2A).

**Figure 2:**
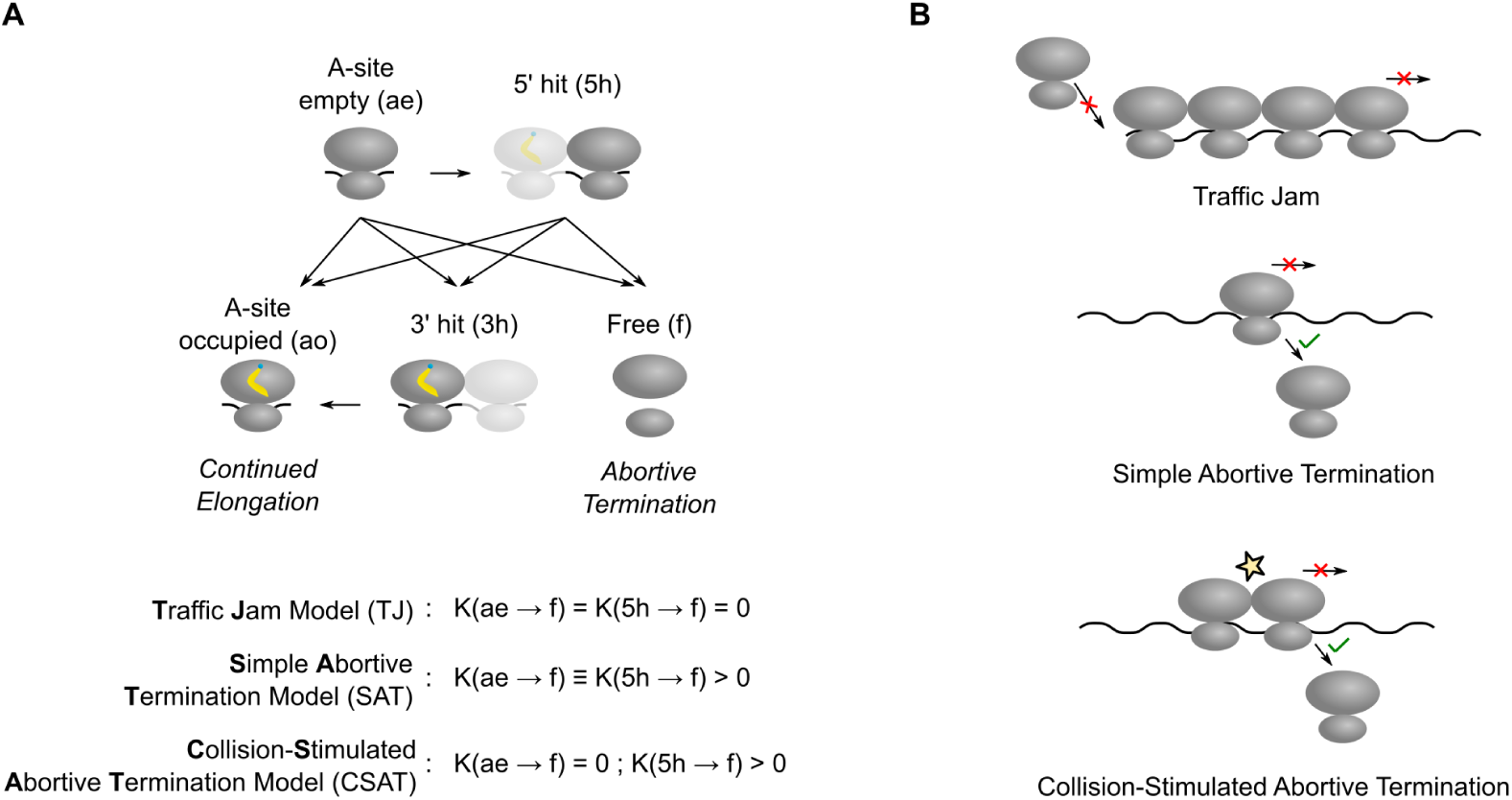
Computational models for ribosome kinetics at stall sites. **A**. Distinct states of ribosome during each elongation cycle that were considered during computational modeling. The three equations indicate the rate of abortive termination from the A-site empty (ae) and the 5’ hit (5h) states in the three different kinetic models considered in this work. **B**. Schematic of the three kinetic models of stalled ribosomes considered in this work.

In our modeling, ribosomes stalled during leucine starvation are represented by the A-site empty state *ae*. Once the leucyl-tRNA is accommodated, A-site empty ribosomes transition to the A-site occupied state *ao*. Ribosomes transition back from the A-site occupied state *ao* to the A-site empty state *ae* upon peptide-bond formation and translocation. Beyond the *ae* and *ao* states, we did not consider additional kinetic states in the normal elongation cycle of the ribosome (19, 21), since such states cannot be resolved using our measurements of protein synthesis rates during leucine starvation. Ribosomes that have dissociated from mRNA either due to normal termination at stop codons or abortive termination at stall sites transition to the free state *f*. Finally, collision between a stalled ribosome with an empty A-site and a trailing ribosome with an occupied A-site results in the stalled ribosome transitioning to the 5’-hit state *5h* and the trailing ribosome transitioning to the 3’-hit state *3h*.

To model ribosome traffic jams, we chose the rate constant for abortive termination of all elongating ribosomes to be zero. Hence if the duration of ribosome stalling is sufficiently long, a queue of trailing ribosomes forms behind the stalled ribosome and ultimately reduces protein synthesis rate by blocking the initiation region. We designate this as the traffic jam (TJ) model (Fig. 2B, top panel).

To model abortive termination, we set the transition rate constant from stalled ribosomes to free ribosomes to be non-zero. Abortive termination occurs selectively at stalled ribosomes, and not at normally elongating ribosomes (31). Even though the mechanistic basis for this selectivity is poorly understood (32, 33), we can account for the selectivity in our modeling by simply setting the abortive termination rate to be zero at all codons except at the stall site (27). We designate this as the simple abortive termination (SAT) model (Fig. 2B, middle panel).

While abortive termination and traffic jams are usually considered as independent molecular processes (34, 35), our definition of kinetic states (Fig. 2A) suggests a more general model in which these processes are coupled. Specifically, we considered a model in which the rate of abortive termination is non-zero only when stalled ribosomes have undergone a collision from a trailing ribosome, i.e. when they are in the *5h* state. We designate this as the collision-stimulated abortive termination (CSAT) model (Fig. 2B, lower panel). As shown below, the CSAT model is closer to experimental measurements of protein synthesis rate than the TJ and SAT models, and it also suggests a mechanistic basis for the selectivity of abortive termination.

### Experimental variables for distinguishing kinetic models of ribosome stalling

Predicting the effect of ribosome stalling on YFP expression in our three kinetic models (Fig. 2B) requires knowledge of the elongation rate and the abortive termination rate of ribosomes at stall-inducing CTA codons during leucine starvation. In principle, these rate constants can be estimated using the ribosome profiling method (27, 36), but sequence-specific and protocol-related biases in ribosome profiling (10, 37, 38) will introduce a large uncertainty in such an estimation. Therefore, we sought to identify experimental variables that would enable us to discriminate between the different kinetic models of ribosome stalling without precise knowledge of the underlying rate constants.

First, we examined the effect of varying the initiation rate of an mRNA with a single stall site in our three kinetic models using stochastic simulations (Fig. 3A, Methods). We chose the elongation and abortive termination rate constants at the stall site such that an mRNA with an initiation rate of 0.3 s^-1^ – a typical value for *E. coli* mRNAs (4, 27) – had the same protein synthesis rate (number of full proteins produced per unit time) in all three models. In the SAT model, varying the initiation rate does not modulate the effect of the stall site on protein synthesis rate (Fig. 3A, blue squares). By contrast, in the TJ and CSAT models, the effect of the stall site on protein synthesis rate was reduced at lower initiation rates (Fig. 3A, green circles and red diamonds). This reduction is more pronounced in the TJ model because, at low initiation rates, ribosome queues do not block the initiation region in the TJ model, while they still lead to collision-stimulated abortive termination in the CSAT model.

**Figure 3:**
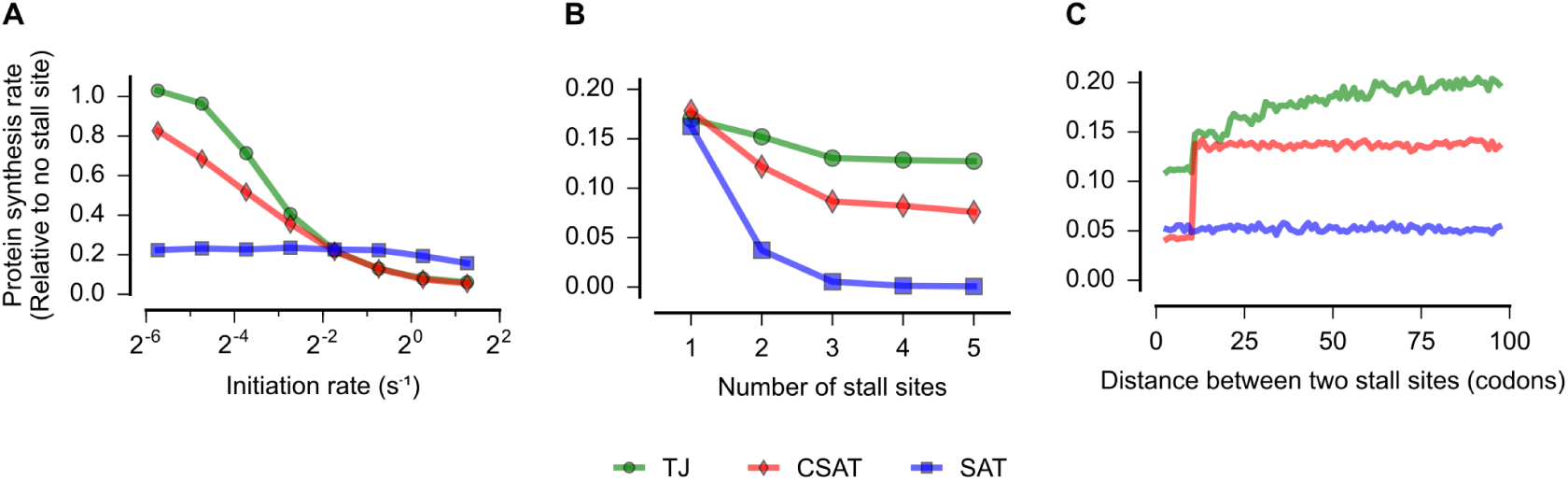
Distinct model predictions upon varying initiation rate, number of stall sites, and distance between stall sites. **A**. Predicted effect of varying the initiation rate of an mRNA with a single stall site on protein synthesis rate. **B**. Predicted effect of varying the number of ribosome stall sites in an mRNA on protein synthesis rate. **C**. Predicted effect of varying the distance between two stall sites in an mRNA on protein synthesis rate. Protein synthesis rate for each mRNA is defined as the number of proteins produced per unit time and is shown relative to an mRNA encoding the same protein, but without stall sites. In A, the duration of stalling in each model was chosen such that the decrease in protein synthesis rate at an initiation rate of 0.3 s^-1^ was equal in the three models. In B and C, the duration of stalling in each model was chosen such that the decrease in protein synthesis rate caused by a single stall site was equal in the three models. K(ae→f) ≡ K(5h→f) = 1 s^-1^ in the SAT model, and K(ae→f) = 0, K(5h→f) = 1 s^-1^ in the CSAT model. Other simulation parameters are in Table S1.

Second, we examined the effect of systematically varying the number of stall sites on an mRNA in our three kinetic models (Fig. 3B). We chose the elongation rate and abortive termination rate constants at stall sites such that the effect of a single stall site on protein synthesis rate was identical between the three models (Table S1). With no further parameter adjustments, we introduced additional identical stall sites such that any two stall sites were separated by at least two ribosome footprints (>60nt). In the traffic jam (TJ) model, additional stall sites had very little effect on protein synthesis rate (Fig. 3B, green circles). In the simple abortive termination (SAT) model, protein synthesis rate decreased exponentially with the number of stall sites (Fig. 3B, blue squares). In the collision-stimulated abortive termination (CSAT) model, the effect of additional stall sites was intermediate between the TJ and SAT models (Fig. 3B, red diamonds). The differential effect of multiple stall sites in the three models can be intuitively understood as follows: In the TJ model, extended queues of ribosomes occur only at the first stall site because the average rate at which ribosomes arrive at subsequent stall sites is limited by the rate at which they elongate past the first stall site. In the CSAT model, ribosome collisions occur at a greater rate at the first stall site, but are not completely prevented at subsequent stall sites due to stochastic ribosome elongation past the first stall site. In the SAT model, abortive termination rate at each stall site does not depend on the presence of other stall sites on the mRNA.

Finally, we considered the effect of varying the distance between two identical stall sites in our kinetic models. In the SAT model, varying the distance between two stall sites does not modulate the effect of the stall site on protein synthesis rate (Fig. 3C, blue). In the CSAT model, when the two stall sites are separated by less than a ribosome footprint, then the frequency of collisions at the stall sites increases, thus resulting in the lower protein synthesis rate in this regime (Fig. 3C, red). In the TJ model, the length of ribosome queues at the first stall site is modulated by the formation of shorter ribosome queues at the second stall site when it within a few ribosome footprints. This interaction results in lower protein synthesis rate when the stall sites are separated by a few ribosome footprints (Fig. 3C, green).

### Measured protein synthesis rates support a collision-stimulated abortive termination model

We tested the predictions from our kinetic models using *yfp* reporters with stall-inducing CTA, CTC and CTT codons during leucine starvation in *E. coli*. First, we measured the effect of varying the initiation rate on the synthesis rate of YFP by mutating either the ATG start codon to near-cognate codons, or by mutating the Shine-Dalgarno sequence (Fig. 4A, inset). We fitted the ribosome elongation rate in the three kinetic models using the measured YFP synthesis rate for the *yfp* variant with the non-mutated initiation region (variant 4 in Fig. 4A), and used this fit to predict the YFP synthesis rate of the other initiation mutants with no remaining free parameters (Methods, Table S2). The effect of a single CTA codon on YFP synthesis rate decreased as the initiation rate of the *yfp* variants was reduced (Fig. 4A, black triangles). Both the TJ and CSAT models predicted the decreasing effect of the CTA codon with lower initiation rate (Fig. 4A, circles & diamonds). By contrast, the predicted YFP synthesis rate from the SAT model was independent of initiation rate (Fig. 4A, squares). This difference between the SAT model and the TJ and CSAT models was also observed for other locations of the CTA codon, or when CTC and CTT codons were introduced into *yfp* (Fig. S1).

**Figure 4:**
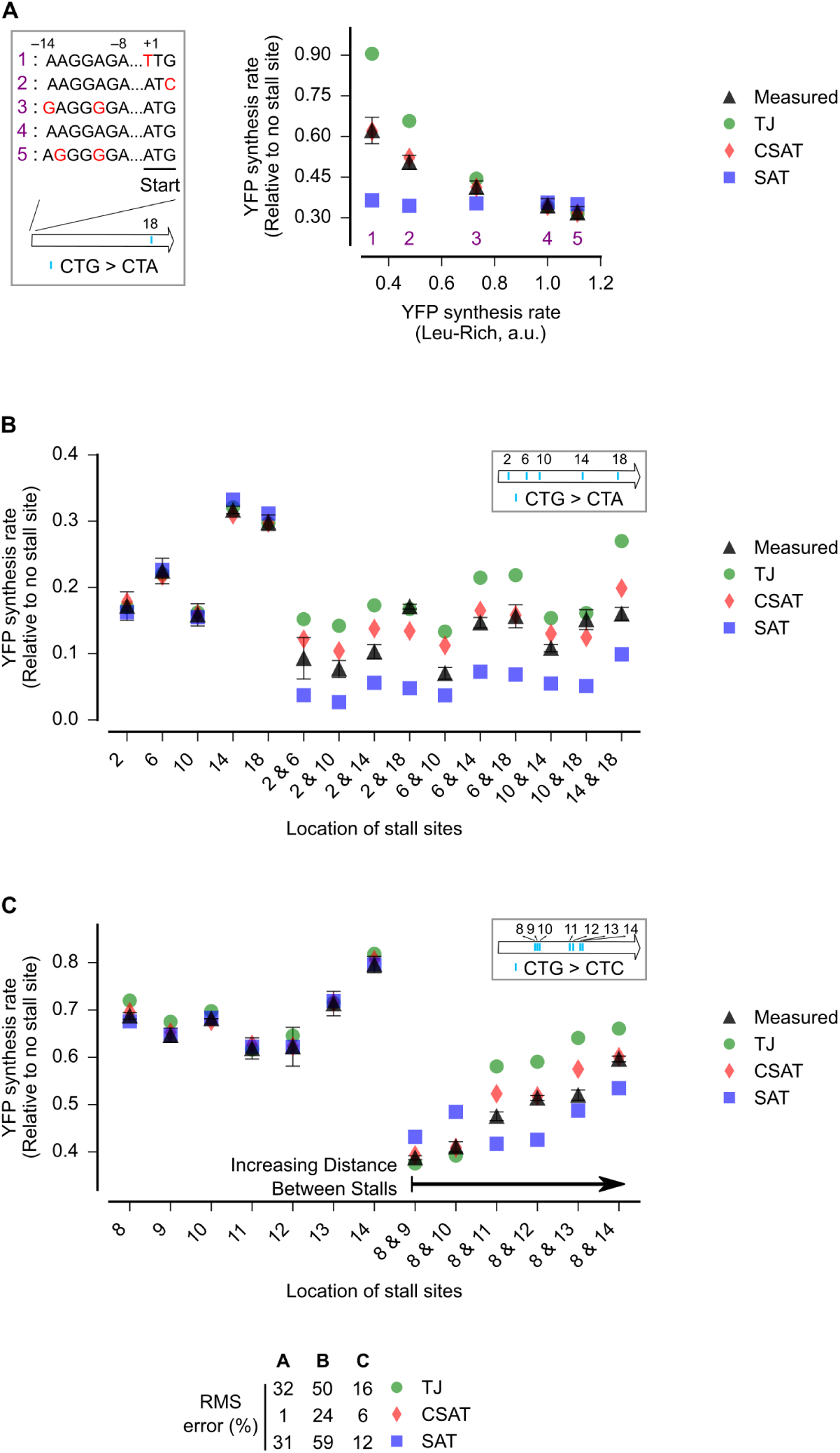
Experimental measurements vs. model prediction for YFP synthesis rate during leucine starvation. **A**. Predicted and measured YFP synthesis rates during Leu starvation from *yfp* reporters with a single CTA codon and one of five initiation region mutations shown in the schematic. The Leu positions are labeled by their order of occurrence along *yfp* (22 Leu codons total). X axis – Measured YFP synthesis rate during Leu rich growth was used as a proxy for the initiation rate of each initiation region mutant. **B**. Predicted and measured YFP synthesis rates during Leu starvation from *yfp* reporters having CTA codons at either one or two among five Leu positions in *yfp* shown in the schematic. X axis - location of CTA codons in each of the 15 *yfp* variants. **C**. YFP synthesis rates and X axis similar to A, but the location of one CTC codon was fixed and the other CTC codon was systematically moved. The Leu codon positions indicated in the schematics in A, B, and C correspond to the following codon positions along *yfp* (with start codon: 1, stop codon: 239): 2: 15, 6: 46, 8: 60, 9: 64, 10: 68, 11: 119, 12: 125, 13: 137, 14: 141, 18: 201. In A, the duration of ribosome stalling in simulations was chosen to reproduce the measured YFP synthesis rate of the reporter with the ‘Init_4’ initiation region. In B and C, the duration of ribosome stalling in simulations was chosen to reproduce the measured YFP synthesis rate of each reporter with single CTA codon. *Bottom legend*: RMS_error_ is the root mean square error between model predictions and measured YFP synthesis rate normalized by the average measured value. RMS_error_ was calculated for mutants 1, 2, 3 and 5 in A, and for the mutants with two CTA/CTC codons in B and C. Error bars indicate standard error over triplicate cultures. Measured and predicted YFP synthesis rate for each *yfp* mRNA is shown relative to a *yfp* mRNA that does not have any CTA/CTC codon. Simulation parameters are shown in Tables S2, S3 and S4.

Second, we tested the effect of multiple stall sites on the synthesis rate of YFP (Fig. 4B). We introduced a single CTA codon at one of five locations among the twenty two leucine codons in *yfp* (Fig. 4B, inset), and we then created all ten pairwise combinations of the five single CTA codons. We used the measured YFP synthesis rate (Fig. 4B, triangles) of the five single CTA variants to fit the ribosome elongation rate at each of the five CTA codon locations in our three kinetic models (Methods, Table S3). These fits, with no remaining free parameters, were used to predict the YFP synthesis rate of the ten double CTA variants during leucine starvation. We found that the TJ model systematically overestimated the YFP synthesis rate for 8 of 10 double CTA variants (Fig. 4B, circles), while the SAT model systematically underestimated the YFP synthesis rate for all 10 double CTA variants during leucine starvation (Fig. 4B, squares). By contrast, the predicted YFP synthesis rates from the CSAT model (Fig. 4B, diamonds) was closer to the measured YFP synthesis rates with less than half the average error of the TJ and CSAT models (Fig. 4, bottom). Similarly, the CSAT model prediction was more accurate when we introduced CTC and CTT codons instead of CTA codons into *yfp* (Fig. S2).

Third, we measured the effect of varying the distance between two stall sites on YFP synthesis rate (Fig. 4C, triangles). We kept the location of a single CTC codon constant and introduced a second CTC codon at increasing distance from the first CTC codon (Fig. 4C, inset). As before, we fitted our three models to the measured YFP synthesis rate of the single CTC variants and used these fits to predict the YFP synthesis rate of the double CTC variants (Methods, Table S4). We found that when the stall sites were separated by more than a few ribosome footprints, the TJ model systematically overestimated the YFP synthesis rate (Fig. 4C, circles) while the SAT model underestimated the YFP synthesis rate (Fig. 4C, squares). When the stall sites were separated by less than a ribosome footprint, the SAT model over-estimated the YFP synthesis rate (Fig. 4C, variants ‘8&9’ and ‘8&10’). By contrast, the CSAT model predictions were close to the measured YFP synthesis rates with no systematic trend as the distance between the two stall sites was varied (Fig. 4C, diamonds). Similarly, the CSAT model was more accurate when a stall site at a different location on YFP was kept constant and the second stall site was moved (Fig. S3).

Collectively, the above experiments reveal that the CSAT model provides a better fit to the measured YFP synthesis rate than either the TJ or the SAT models when the number of stall sites, the distance between stall sites, and the initiation rate are systematically varied during leucine starvation in *E. coli*.

### Selectivity, robustness and ribosome density in the collision-stimulated abortive termination model

The ability of the CSAT model to account for YFP synthesis rates from our reporters led us to examine whether this model is consistent with other expected features of ribosome stalling during amino acid starvation in *E. coli*.

First, only a small fraction of ribosomes are expected to prematurely terminate from mRNAs without stall sites (27, 39, 40). Consistent with this expectation, predicted protein synthesis rates from reporters without stall sites did not decrease when the abortive termination rate was increased in the CSAT model (Fig. 5A, red). This selectivity towards stalled ribosomes naturally arises in the CSAT model from the requirement for ribosome collisions to cause abortive termination. By contrast, protein synthesis rates from reporters without stall sites decreased with increasing abortive termination rate in a SAT model in which abortive termination was not explicitly specified to be selective for stalled ribosomes (Fig. 5A, blue vs. pink).

**Figure 5:**
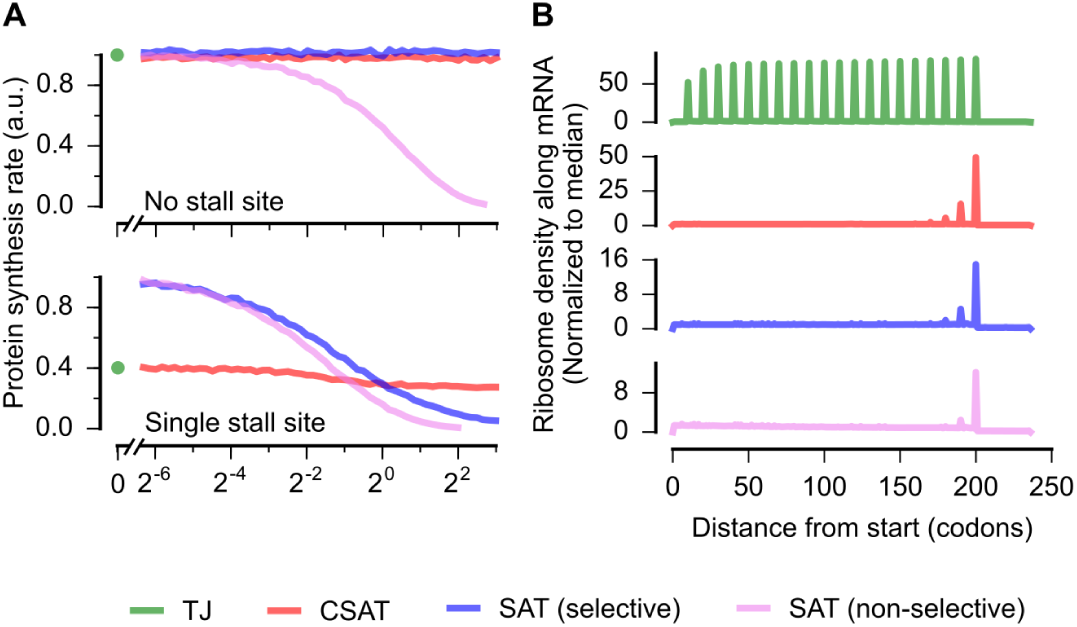
Model predictions for the effect of varying abortive termination rate on protein synthesis rate and ribosome occupancy. **A**. Predicted effect of varying the abortive termination rate on protein synthesis rate from a *yfp* mRNA having no stall site (top panel) or a single stall site (bottom panel). The TJ model corresponds to an abortive termination rate of zero, and is shown as a single point at the left. The selective SAT model has non-zero abortive termination rate at only the codon corresponding to the stall site. The non-selective SAT model has non-zero abortive termination rate at all codons along the *yfp* mRNA. Overlapping curves for CSAT and SAT (selective) models in top panel were manually offset for clarity. **B**. Predicted ribosome occupancy on a *yfp* mRNA with a single stall site at the 200^th^ codon. Ribosome occupancy is normalized by its median value across the mRNA. Other simulation parameters shown in Table S5.

Second, the frequency of abortive termination is known to be robust to over-expression of factors that rescue stalled ribosomes (41). Consistent with this observation, we found that increasing the abortive termination rate in the CSAT model predicted only a minor effect on protein synthesis rate from an mRNA with a single stall site (Fig. 5B, red). By contrast, in both the selective and non-selective SAT models, protein synthesis rate from an mRNA with a single stall site continuously decreased as the abortive termination rate was increased (Fig. 5B, blue and pink). The robustness of the CSAT model to varying abortive termination rates arises because the frequency of ribosome collisions limit the actual rate of abortive termination at stall sites.

Finally, previous ribosome profiling measurements have detected a queue of only a few ribosomes at CTA codons during leucine starvation in *E. coli* (27). Consistent with this observation, the length of ribosome queues at the stall site predicted by the CSAT model is limited to a few ribosomes even when the stall site is located ~200 codons from the start codon (Fig. 5C, red). A similar queue of few ribosomes is also observed in the SAT model (Fig. 5C, blue and pink). By contrast, the TJ model predicts a queue of around 20 ribosomes when stall sites are located ~200 codons from the start codon (Fig. 5C, green).

## Discussion

In this work, we used a combination of computational modeling and reporter-based measurements of protein synthesis rate to constrain ribosome kinetics at stall sites during leucine starvation in *E. coli*. Our approach allowed us to quantitatively test two previously proposed models for how ribosome stalling decreases protein expression, namely ribosome traffic jams that block initiation (TJ model) and simple abortive termination of stalled ribosomes (SAT model). We also considered a novel model in which ribosome collisions stimulate abortive termination of stalled ribosomes (CSAT model). Our integrated approach allowed us to infer the extent to which each of these three kinetic models accounted for the measured protein synthesis rate from a library of *yfp* variants.

The TJ model has been considered theoretically in several studies (8, 11, 42). While queues of ~7 ribosomes have been detected *in vitro* (43), ribosome profiling studies have revealed a queue of only a few ribosomes at stall sites *in vivo* (10, 27, 44). These smaller queues can modulate protein expression only if the stall site is within a few ribosome footprints from the start codon (11, 45, 46). Nevertheless, recent studies on EF-P dependent pauses in bacteria and rare-codon dependent pauses in yeast suggest that the TJ model underlies the decreased protein expression when stall sites are over 100 codons away from the start codon (9, 12). These conclusions were based on observations that decreasing initiation rate of ribosomes on reporters reduced the effect of stall sites on protein expression (9, 12). This modulatory effect of initiation rate was also observed in our experiments (Fig. 4A). However, we find that both the TJ and CSAT models predict this modulatory effect of initiation rate (Fig. 3A), while only the CSAT model predicts a queue of few ribosomes (Fig. 5C) that is observed experimentally. Thus, collision-stimulated abortive termination is a plausible alternative mechanism to the traffic jam models proposed in previous studies of ribosome stalling (9, 12).

Simple kinetic partitioning between normal elongation and abortive termination has been proposed as a possible mechanism for how ribosome rescue factors might act at ribosomes that are stalled within an mRNA (47, 48). However, our modeling indicates that this non-selective mechanism of abortive termination will result in decreased protein expression from mRNAs that do not have stall sites (Fig. 5A). This observation can be intuitively understood from the fact that even a small probability of abortive termination during each elongation cycle will be multiplicatively amplified over the course of translating a typical *E. coli* protein with 300 amino acid residues.

We found that the CSAT model is consistent with measured protein synthesis rates and ribosome density near stall sites during amino acid starvation in *E. coli*. Nevertheless, the exact mechanism of abortive termination that is stimulated by ribosome collisions remains to be established. Specifically, ribosome collisions could either stimulate spontaneous drop-off of stalled ribosomes, or they could stimulate the activity of quality control pathways such as the tmRNA and the ArfA systems that rescue stalled ribosomes (13, 49). In the latter case, ribosome collisions might allow the quality control pathway to selectively recognize ribosomes that have been stalled for an extended duration over ribosomes that are transiently stalled due to the stochasticity of normal elongation (33). In this sense, the frequency of ribosome collisions can provide a natural timer for achieving selectivity of quality control pathways towards stalled ribosomes (48). Further, the robustness of the CSAT model to changes in the abortive termination rate (Fig. 5B) can buffer against cell-to-cell variation in the concentration of quality control factors that mediate abortive termination.

Despite the better fit provided by the CSAT model to our measured YFP synthesis rates, there still remains a residual error in its prediction (Fig. 4, bottom). This error might arise from several simplifying assumptions in our definition of the CSAT model, which we made in order to emphasize its qualitative difference with the TJ and SAT models (Fig. 2A). Specifically, we assumed the rate of abortive termination to be zero in the absence of ribosome collisions. Relaxing this assumption is likely to provide a better fit to our measurements, but it will introduce extra free parameters while not offering additional mechanistic insight into the kinetics of abortive termination. We also assumed the rate of abortive termination to be zero for the trailing ribosome in the collided state (*3h* in Fig. 2A), since there is no biochemical evidence for such a process. These assumptions could be relaxed based on evidence from future biochemical and structural studies of ribosome queues formed at stall sites.

The inverse approach used in our work relied on model predictions that did not depend sensitively on underlying kinetic parameters such as the elongation rate and the abortive termination rate at stall sites. By the same token, our work cannot be used to infer the exact values of these kinetic parameters *in vivo*. In the future, these parameters might be directly accessible using recent imaging approaches for measuring translation kinetics on single mRNPs (3).

## Data Accession

Raw data and programming code for reproducing all figures in this paper is publicly available at: http://github.com/rasilab/ferrin_1.

## Acknowledgements

We thank Robert Bradley, Allen Buskirk, Premal Shah, Hani Zaher and Brian Zid for discussions. Funding for this work was provided by NIH grant R35 GM119835, NIH grant R00 GM107113, and startup funds from the Fred Hutchinson Cancer Research Center. The computations in this paper were run on the Gizmo cluster supported by the Scientific Computing group at the Fred Hutchinson Cancer Research Center.

## Contributions

M.F and A.R.S. designed research, analyzed data, and wrote the manuscript. M.F performed experiments. A.R.S performed computational modeling.

---

**Figure S1:**
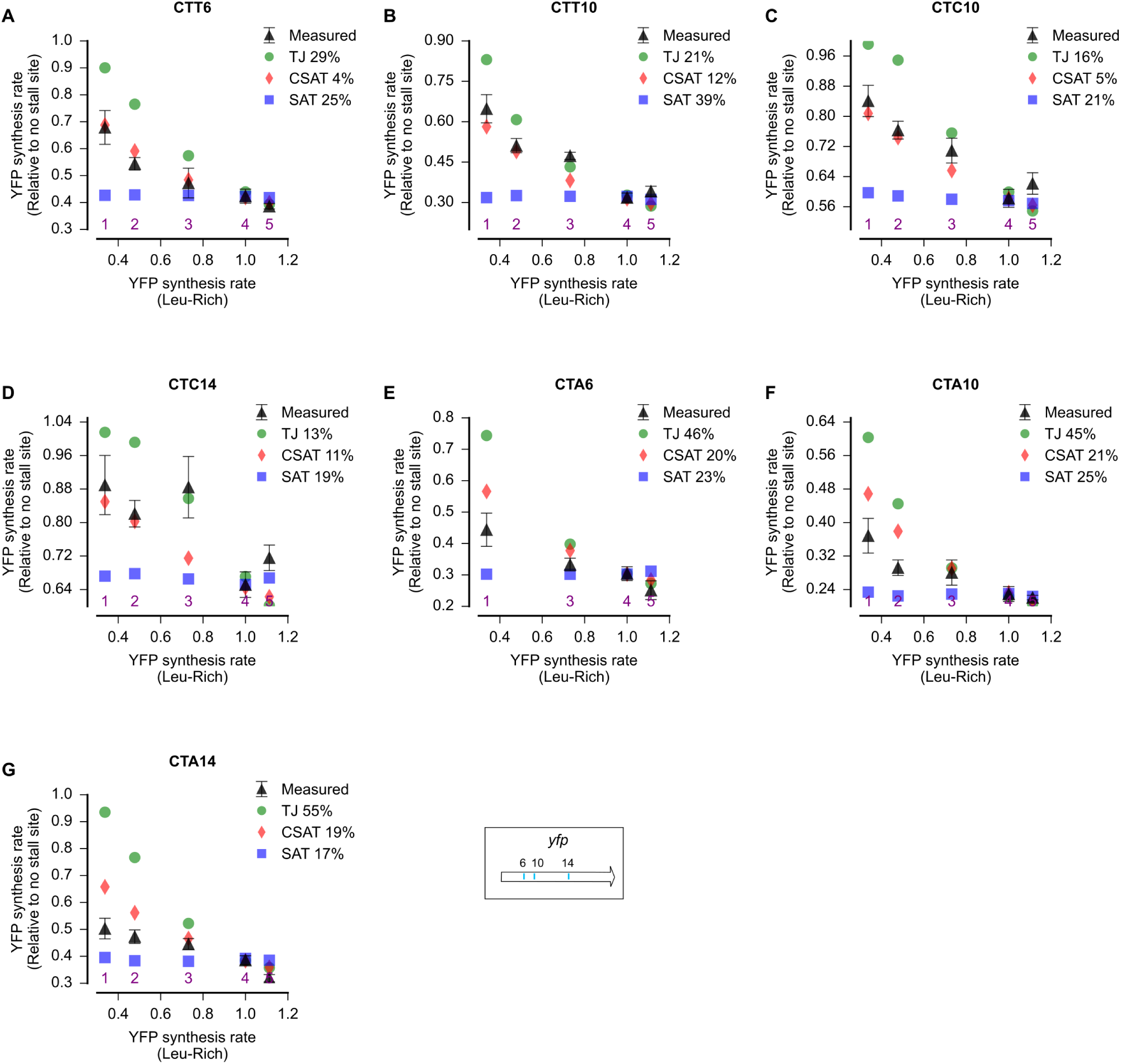
Effect of initiation rate variation on predicted and measured YFP synthesis rates (Related to Fig. 4A) Predicted and measured YFP synthesis rates during Leu starvation from *yfp* reporters with a single stall-inducing codon and one of five initiation region variants in Fig. 4A (labeled 1–5). X axis – Measured YFP synthesis rate during Leu rich growth was used as a proxy for the initiation rate of each initiation region mutant. Each panel represents data for initiation rate variants having the indicated stall-inducing codon (CTT, CTC or CTA) at one of three different leucine codon locations along *yfp*. The Leu positions are labeled by their order of occurrence along *yfp* (22 Leu codons total). The location of the 6^th^,10^th^ and 14^th^ Leu codons along *yfp* are shown in the schematic at the bottom. The numerical values next to the legend indicate the RMS_error_ between measured and predicted YFP synthesis rates from each model, and was calculated similar to Fig. 4A. All simulation parameters are shown in Table S2.

**Figure S2:**
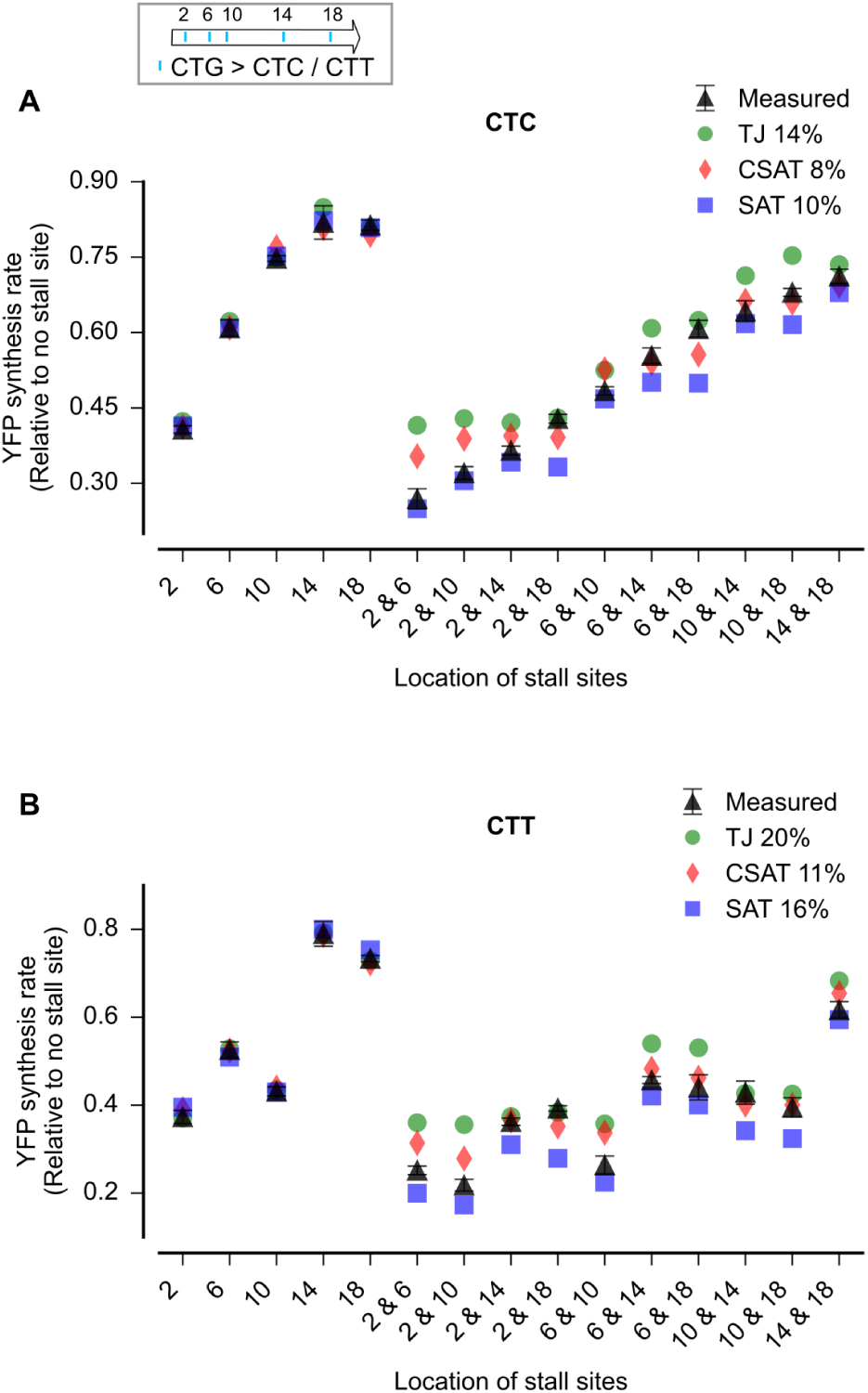
Effect of multiple stall sites on predicted and measured YFP synthesis rates (Related to Fig. 4B) Predicted and measured YFP synthesis rates during Leu starvation from *yfp* reporters having CTC (A) or CTT (B) codons at either one or two among five Leu positions in *yfp* shown in the schematic. The Leu positions are labeled by their order of occurrence along *yfp* (22 Leu codons total). X axis - location of CTC/CTT codons in each of the *yfp* variants. The numerical values next to the legend indicate the RMS_error_ between measured and predicted YFP synthesis rates from each model, and was calculated similar to Fig. 4B. All simulation parameters are shown in Table S3.

**Figure S3:**
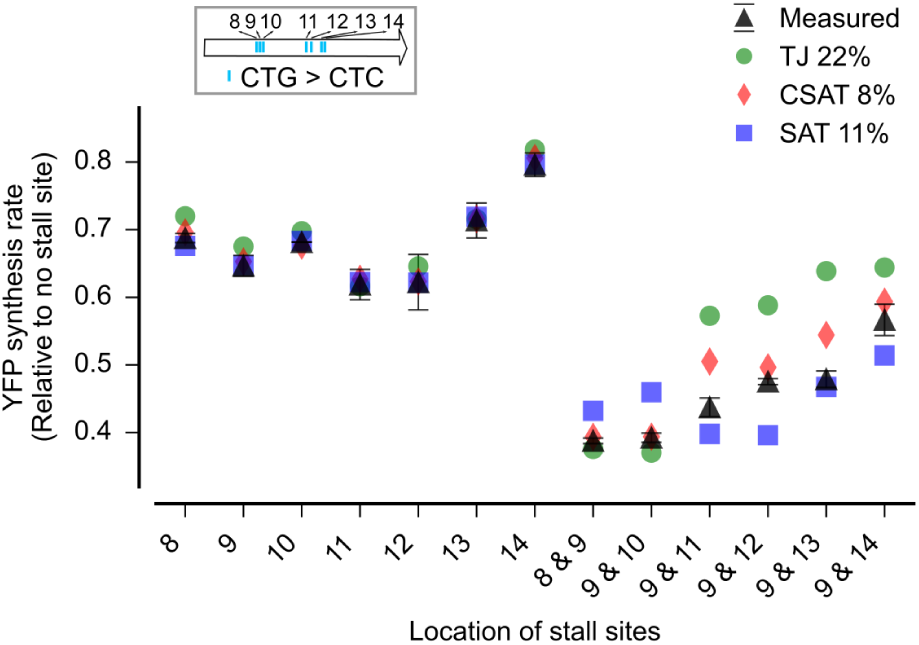
Effect of distance between stall sites on predicted and measured YFP synthesis rates (Related to Fig. 4C) Predicted and measured YFP synthesis rates during Leu starvation from *yfp* reporters in which the location of one CTC codon was fixed and the other CTC codon was systematically moved. The Leu positions are labeled by their order of occurrence along *yfp* (22 Leu codons total) and their approximate positions along *yfp* are indicated in the schematic. X axis - location of CTC codons in each of the *yfp* variants. The numerical values next to the legend indicate the RMS_error_ between measured and predicted YFP synthesis rates from each model, and was calculated identical to Fig. 4C. All simulation parameters are shown in Table S4.

## Materials and Methods

### Bacterial strains and plasmids

All experiments in this study were performed using an *E. coli* strain (*28*) that is auxotrophic for leucine and containing the *tet* repressor gene for inducible control of reporter genes (ecMF1). All fluorescent reporters in this study were cloned into a very low copy expression vector (SC*101 *ori*, 3-4 copies per cell) used in our previous work (*28*) (pASEC1, Addgene plasmid #53241). The fluorescent reporters genes were based off a yellow fluorescent protein sequence (*yfp0*) present in pASEC1, which encodes a fast-maturing ‘Venus’ variant of YFP. All 22 leucine codons in *yfp0* were chosen as CTG. For constructing *yfp* reporters with single stall sites, the corresponding CTG codon in *yfp0* was mutated to CTA, CTC, or CTT by encoding these mutations in oligos and using Gibson assembly (*50*). A *yfp* variant with 7 leucine codons mutated to CTA was used in all plate reader experiments as a control for the lower limit of detection of YFP fluorescence under Leu starvation (ecMF112). Variants of *yfp* with two CTA, CTC or CTT codons were constructed by Gibson assembly of PCR fragments from the corresponding single codon variants of *yfp*. The start codon and Shine-Dalgarno sequence variants of *yfp* were generated by encoding these mutations in one of the PCR oligos for *yfp.* The *3xflag-yfp* variants were generated by the addition of a 22 codon sequence at the 5’ end that encoded a 3X-FLAG peptide used in our previous work (*28*). All strains and plasmids used in this study are listed in Table S7. Strains are available upon request.

### Growth and fluorescence measurements

Overnight cultures were inoculated in biological triplicates from freshly grown single colonies or patched colonies from glycerol stocks. Overnight cultures were grown in a modified MOPS rich defined medium (*28*, *51*) made with the following recipe: 10X MOPS rich buffer, 10X ACGU nucleobase stock, 100X 0.132M K_2_HPO_4_ were used at 1X final concentration as in the original recipe. In addition, the overnight growth medium contained 0.5% glucose as carbon source and 800µM of 19 amino acids and 10mM of serine. pH was adjusted to 7.4 using 1M NaOH and appropriate selective antibiotic (50µg/ml carbenicillin) was added. 200ng/ml of anhydro-tetracycline (aTc) was also added in order to induce the pLtetO1 promoter. 1ml overnight cultures were grown in 2ml deep 96-well plates (AB0932, Fisher) at 30℃ with shaking at 1200rpm (Titramax 100 shaker) for 12 to 16 hours.

For leucine starvation time course experiments, overnight cultures were diluted 1:100 into 150µl of the same MOPS rich-defined medium as the overnight cultures. However leucine was added at 100µM and supplemented with its methyl ester analog at 160µM (AC125130250, Fisher). Addition of the leucine methyl ester results in a steady but limiting supply of leucine due to slow hydrolysis of the ester, and this enables extended and accurate measurements of protein synthesis rate under Leu starvation condition (*28*). Except for leucine, the remaining 19 amino acids were present at the overnight culture concentrations during the leucine starvation experiments.

Diluted overnight cultures were grown in 96-well plates (3595, Costar) at 30℃ with shaking at 1200rpm (Titramax 100 shaker). A 96-well plate reader (Infinite M1000 PRO, Tecan) was used to monitor cell density (absorbance at 600nm) and YFP synthesis (fluorescence, excitation 504nm and emission 540nm). Each plate was read every 15 min and shaken in between readings for a total period of 6–10 hours.

For experiments in Fig. 1, overnight cultures were grown without aTc and diluted 1:1000 into the same medium. Then when the OD_600_ reached 0.5, the cells were spun down at 3000g for 5 min and then re-suspended in the same medium, but either with or without leucine, and with aTc for reporter induction. Fluorescence, Westerns, qRT-PCR measurements in Fig. 1 were performed from these cultures after shaking at 37℃, 200rpm for 20 min with leucine or 60 min without leucine.

### Polysome profiling

Overnight cultures were diluted 1:200 into 400ml MOPS rich defined medium and grown at 37℃ to an OD_600_ of 0.2. Cells were harvested by vacuum filtration on a 0.2 μm nitrocellulose membrane (BA83, GE) and subsequently cut in half. One half was added to 200ml MOPS rich defined medium, the other to 200ml of same medium but without leucine. After growth at 37℃ for either 20 minutes (Leu-rich cultures) or 1 hour (Leu starvation cultures), cells were harvested by vacuum filtration again. Cells were scraped from the membrane using a plastic spatula before the membrane became totally dry, and then immediately submerged in liquid nitrogen and stored at -80°C. Frozen cells were then resuspended in 0.7 ml bacterial lysis buffer (20 mM Tris pH 8.0, 10 mM MgCl2, 100 mM NH4Cl, 2 mM DTT, 0.1% NP‐40, 0.4% Triton X‐100, 100 U/mL DNase I, and 1 mM chloramphenicol) and lysed using glass beads (G1277, Sigma) by vortexing 4×30s at 4℃ with 60s cooling on ice in between. The lysate was clarified by centrifugation at 21,000g for 10 min and supernatant was transferred to a fresh tube.

Lysate RNA concentration was quantified by A_260_ (Thermo Scientific Nanodrop) and 100–200µl of lysate containing 0.5 mg RNA was loaded onto a 10-50% sucrose gradient made with 20 mM Tris pH 8.0, 10 mM MgCl2, 100 mM NH4Cl, and 2 mM DTT. Polysomes were separated by centrifugation in an SW41 rotor at 35,000 rpm for 3 h at 4°C. Gradients were then fractionated into 15 fractions containing 25.6 ng spike-in control FLuc mRNA. RNA from each fraction was column-purified (Quick-RNA Miniprep, Zymo) along with in-column DNase I digestion (Zymo).

### Total RNA extraction

Phenol-chloroform extraction method was used to obtain total RNA. 10ml of cells were quickly chilled in an ice water bath and harvested by centrifugation at 3000g for 5 min. Cell pellets were re-suspended in 500µl of 0.3M sodium acetate, 10mM EDTA, pH 4.5. Resuspended cells were mixed with 500µl of acetate-saturated phenol-chloroform at pH 4.5 and 500µl of acid-washed glass beads (G1277, Sigma). The mixture was shaken in a vortexer for 3 min at 4℃ and then clarified by centrifugation at 21,000g for 10 min. The aqueous layer was extracted twice with acetate-saturated phenol-chloroform at pH 4.5 and once with chloroform. Total RNA was precipitated with an equal volume of isopropanol and washed with 70% ethanol and finally re-suspended in 200µl of RNase-free 10mM Tris pH 7.0. 200ng of the total RNA was treated with DNase (M0303S, NEB) to remove residual DNA contamination (manufacturer’s instructions were followed). The DNA-free RNA was column-purified (Quick-RNA Miniprep, Zymo).

### Reverse transcription and quantitative PCR

Reverse transcription (RT) was performed using 10–20ng of DNA-free RNA and Maxima reverse transcriptase (EP0741, Thermo), used according to manufacturer’s instructions. Random hexamer primers were used for priming the RT reaction. At the end of the RT reaction, the 10µl RT reaction was diluted 20-fold and 5µl of this diluted sample was used as template for qPCR in the next step. qPCR was performed using Maxima SYBR Green/ROX qPCR Master Mix (FERK0221, Thermo) and manufacturer’s instructions were followed. qPCR was performed in triplicates for each RT reaction and appropriate negative RT controls were used to confirm the absence of DNA contamination. *gapA* mRNA was used as internal reference to normalize all other mRNA levels. Primers for qPCR (Table S5) were from our previous work (*28*). ∆Ct method was used to obtain relative mRNA levels. Analysis was implemented using Python 2.7 libraries. Code for analysis and plotting of figures starting from raw qPCR data is publicly available at http://github.com/rasilab/ferrin_1 as Jupyter notebooks (*52*).

### Western blotting

Cells were harvested by centrifugation and protein was precipitated by mixing TCA to a final concentration of 10%. The mixture was incubated on ice for 15 min and the supernatant removed. Protein pellets were re-suspended in 100 μl 1X Laemmli Buffer (Biorad), boiled at 99℃ for 5min, and then loaded onto each lane of a 4–20% polyacrylamide gel (Biorad) and SDS-PAGE was carried out at 200V for 50 min. Proteins were transferred to a nitrocellulose membrane at 500mA for 60 min using a wet-transfer apparatus (Biorad). The membrane cut along the 50kD marker and both halves were blocked in Odyssey PBS Blocking Buffer (Li-cor) for 60 min. The lower-MW half was incubated with a 1:6000 dilution of an anti-FLAG antibody (F3165, Sigma), and the higher-MW half in the same dilution of an anti-sigma 70 antibody (WP004, Neoclone), each in 15ml of Odyssey PBS Blocking Buffer with shaking at 4℃ overnight. After washing 4×5 min with TBST, the membrane was incubated with 1:10,000 dilution of a secondary dye-conjugated antibody (925-68072, Li-cor) in 15ml of Odyssey PBS Blocking Buffer with shaking at room temperature for 60 min. After washing 4×5 min with PBS, the membrane was imaged using a laser-based fluorescence imager.

### Growth and fluorescence data analysis

OD_600_ and YFP fluorescence were recorded as time series for each well of a 96-well plate. Background values for OD_600_ and YFP fluorescence were subtracted based on measurements from a well with just growth medium. Time points corresponding to Leu-rich growth and Leu starvation were identified by manual inspection of OD_600_ curves. The onset time of starvation was automatically identified as the time point at which YFP/ OD_600_ reached a minimum value. YFP synthesis rate during Leu-rich exponential growth was defined as the average of YFP/ OD_600_ values for the 3 points around the onset time of starvation. YFP synthesis rate during Leu starvation was defined as the slope of a linear fit to the fluorescence time series in the Leu starvation regime. YFP synthesis rates for individual wells were averaged over biological replicate wells for calculation of mean and standard error. Analysis was implemented using Python 2.7 libraries. Code for analysis and plotting of figures starting from raw plate reader data is publicly available at http://github.com/rasilab/ferrin_1 as Jupyter notebooks (*52*).

### Simulation

The kinetic models in Fig. 2 were implemented as stochastic simulations in the C++ object-oriented programming language. Separate classes were defined to represent ribosomes, mRNA transcripts, gene sequences, tRNAs and codons. Each elongating ribosome was represented as an instance of the *Ribosome* class. The four distinct states of the elongating ribosome (*ae*, *ao*, *5h*, *3h* in Fig. 2A) were tracked using three *bool* properties of the *Ribosome* class: *AsiteEmpty*, *hitFrom5Prime*, and *hitFrom3Prime*. The identities of the tRNAs occupying the A-site and P-site of the elongating ribosome were tracked. Only the aggregate number of ribosomes in the free state (*f* in Fig. 2A) was tracked. Instances of the *transcript* class were used to track the number of proteins produced from each transcript. The *gene*, *tRNA* and *codon* classes were used as data structures and their properties did not change during the course of the simulation.

Each simulation considered translation of two mRNA molecules that both encoded YFP. The first mRNA molecule was a control *yfp* sequence without any CTA, CTC or CTT codon. The second mRNA molecule was the test *yfp* sequence with the CTA, CTC or CTT codon as specified for individual simulations.

We simulated four different molecular processes during translation: initiation, elongation, aminoacylation and abortive termination. The rates of all other steps in translation such as termination and ribosome recycling were set to be instantaneous.

The initiation rate of all mRNA sequences was set as 0.3s^-1^ [a typical value for *E. coli* mRNAs (*4*, *27*)] except when this rate was explicitly varied, either to demonstrate its effect in our kinetic models (Fig. 3A) or for experimental fits (Fig. 4A, Fig. S1). For the experimental fits in Fig. 4A and S1, the measured YFP synthesis rate of the initiation region mutants during Leu-rich growth relative to the starting sequence (Init_4 in Fig. 4A) was used to scale the default initiation rate of 0.3s^-1^.

Elongation cycle of ribosomes at each codon was divided into two steps:

In the first elongation step, the cognate tRNA is accommodated into the A-site. The rate of tRNA accommodation was chosen to be non-zero only when ribosomes are in the *ae* state. The tRNA accommodation rate for all codons was calculated as the product of a pseudo first-order rate constant (2×10^7^ M^-1^s^-1^), the concentration of individual tRNAs, and a weight factor to account for codon-anticodon pairing strength. The concentration of tRNAs and the weight factors were based on measured concentration of *E. coli* tRNAs (*53*) and known wobble-pairing rules (*27*, *36*). Leucine starvation was simulated using a previous whole cell model of translation (*27*). The steady-state charged fraction of all tRNAs from this whole-cell model during leucine starvation was used for our *yfp* reporter simulation as the default values. To fit the measured YFP synthesis rate of single stall-site variants (Fig. 4B, 4C, S2, S3), the tRNA accommodation rate at CTA, CTC and CTT codons was systematically varied in the three kinetic models. These fit values for used for illustrating the predictions from the kinetic models in Fig. 3 and Fig. 5.

In the second elongation step, peptide bond is formed and ribosomes translocate to the next codon. This rate was set to be 22s^-1^ and equal to the maximum measured rate of *in vivo* elongation (*54*).

The aminoacylation rate for all tRNAs was calculated as the product of a pseudo first-order rate constant (2×10^10^ M^-1^s^-1^) and the concentration of individual tRNAs. Even though we simulated this process explicitly, we did not lower this rate for leucine tRNAs to simulate leucine starvation; Instead, we accounted for leucine starvation by using the steady-state charged fraction of leucine tRNAs from our whole-cell model as mentioned above in our discussion of tRNA elongation rate. This modified procedure enabled us to simulate the translation of just the *yfp* reporters without considering all the endogenous mRNAs in the cell.

The abortive termination rate was set to a value of 1s^-1^ in the SAT and CSAT models and 0s^-1^ in the TJ model, except when this rate was explicitly varied (Fig. 5A). We chose this rate to be of the same approximate value as in our ribosome profiling studies (*27*). The exact value of this rate is not critical in our SAT and CSAT models since the fitted value of the elongation rate gets adjusted to fit the measured protein synthesis rate from our YFP reporters.

The simulations used a stochastic Gillespie algorithm that was implemented in earlier studies (*27*, *36*). Each simulation was run until 10,000 full-length YFP molecules were produced from the first *yfp* mRNA without CTA, CTC or CTT codons. The number of full-length YFP molecules produced from the second *yfp* mRNA with stall sites in the same duration was used to calculate the YFP synthesis rate (in Fig. 3,4,5,S1,S2,S3) after normalizing by 10,000. Time-averaged ribosome density on each mRNA was also tracked during the simulation run after 100 YFP molecules were produced from the first *yfp* mRNA, and this density was median-normalized for plotting in Fig. 5B.

Code for creating simulation input files, running the simulation, and plotting of figures starting from simulation results is publicly available at http://github.com/rasilab/ferrin_1 as Jupyter notebooks (*52*). Parameters common to all simulations are listed in Table S6. Parameters specific to simulations in individual figures are listed in Tables S1–S5.

## Supplementary Tables

**Table S1:** Simulation parameters for Fig. 3.

| Parameter | Value | Note |
| --- | --- | --- |
| Stall site identity | CTA | Panel A, B, C |
| Stall site location (codon number along *yfp*) | 46 | Panel A |
| Initiation rate | [0.019, 0.038, 0.075, 0.15, 0.3, 0.6, 1.2, 2.4] s<sup>-1</sup> | Panel A |
| Stall site locations (codon number along *yfp*) | 15, 46, 68, 141, 201 | Panel B, stall sites were added from 5' to 3'. |
| First stall site location (codon number along *yfp*) | 60 | Panel C |
| Second stall site locations (codon number along *yfp*) | 63, 64, ... 238 | Panel C |
| tRNA accommodation rate at stall site (TJ model) | 0.065s<sup>-1</sup> | Panels A, B, C |
| tRNA accommodation rate at stall site (SAT model) | 0.286s<sup>-1</sup> | Panels A, B, C |
| tRNA accommodation rate at stall site (CSAT model) | 0.085s<sup>-1</sup> | Panels A, B, C |

The tRNA accommodation rate at stall sites (which sets the duration of stalling) were chosen as described in the main text and figure caption. All other parameters have values shown in Table S6.

**Table S2:** Simulation parameters for Fig. 4A and Fig. S1.

| Parameter | Value | Note |
| --- | --- | --- |
| Stall site identity | CTA, CTC, CTT | Fig. 4A, S1 |
| Stall site locations as codon number along *yfp*<br>(leucine codon number as indicated along X axis) | 46 (6), 68 (10), 141 (14), 201 (18) | Different sites were chosen as indicated in panel title (Fig. S1) and schematic. |
| tRNA accommodation rate at CTA68 (CSAT model) | 0.0919s<sup>-1</sup> | Fit from experiment |
| tRNA accommodation rate at CTA141 (CSAT model) | 0.178s<sup>-1</sup> | Fit from experiment |
| tRNA accommodation rate at CTA201 (CSAT model) | 0.152s<sup>-1</sup> | Fit from experiment |
| tRNA accommodation rate at CTA46 (CSAT model) | 0.132s<sup>-1</sup> | Fit from experiment |
| tRNA accommodation rate at CTC68 (CSAT model) | 0.354s<sup>-1</sup> | Fit from experiment |
| tRNA accommodation rate at CTC141 (CSAT model) | 0.433s<sup>-1</sup> | Fit from experiment |
| tRNA accommodation rate at CTT68 (CSAT model) | 0.137s<sup>-1</sup> | Fit from experiment |
| tRNA accommodation rate at CTT46 (CSAT model) | 0.205s<sup>-1</sup> | Fit from experiment |
| tRNA accommodation rate at CTA68 (SAT model) | 0.296s<sup>-1</sup> | Fit from experiment |
| tRNA accommodation rate at CTA141 (SAT model) | 0.638s<sup>-1</sup> | Fit from experiment |
| tRNA accommodation rate at CTA201 (SAT model) | 0.543s<sup>-1</sup> | Fit from experiment |
| tRNA accommodation rate at CTA46 (SAT model) | 0.442s<sup>-1</sup> | Fit from experiment |
| tRNA accommodation rate at CTC68 (SAT model) | 1.41s<sup>-1</sup> | Fit from experiment |
| tRNA accommodation rate at CTC141 (SAT model) | 1.96s<sup>-1</sup> | Fit from experiment |
| tRNA accommodation rate at CTT68 (SAT model) | 0.484s<sup>-1</sup> | Fit from experiment |
| tRNA accommodation rate at CTT46 (SAT model) | 0.73s<sup>-1</sup> | Fit from experiment |
| tRNA accommodation rate at CTA68 (TJ model) | 0.0684s<sup>-1</sup> | Fit from experiment |
| tRNA accommodation rate at CTA141 (TJ model) | 0.119s<sup>-1</sup> | Fit from experiment |
| tRNA accommodation rate at CTA201 (TJ model) | 0.106s<sup>-1</sup> | Fit from experiment |
| tRNA accommodation rate at CTA46 (TJ model) | 0.0948s<sup>-1</sup> | Fit from experiment |
| tRNA accommodation rate at CTC68 (TJ model) | 0.193s<sup>-1</sup> | Fit from experiment |
| tRNA accommodation rate at CTC141 (TJ model) | 0.22s<sup>-1</sup> | Fit from experiment |
| tRNA accommodation rate at CTT68 (TJ model) | 0.0975s<sup>-1</sup> | Fit from experiment |
| tRNA accommodation rate at CTT46 (TJ model) | 0.137s<sup>-1</sup> | Fit from experiment |
| Initiation rate of Init_1 variants | 0.101s<sup>-1</sup> | Fit from experiment |
| Initiation rate of Init_2 variants | 0.143s<sup>-1</sup> | Fit from experiment |
| Initiation rate of Init_3 variants | 0.219s<sup>-1</sup> | Fit from experiment |
| Initiation rate of Init_4 variants | 0.3s<sup>-1</sup> | Default value |
| Initiation rate of Init_5 variants | 0.334s<sup>-1</sup> | Fit from experiment |

All other parameters have values shown in Table S6.

**Table S3:** Simulation parameters for Fig. 4B and Fig. S2.

| Parameter | Value | Note |
| --- | --- | --- |
| Stall site identity | CTA | Fig. 4B |
| Stall site identity | CTC | Fig. S2A |
| Stall site identity | CTT | Fig. S2B |
| Stall site locations as codon number along *yfp*<br>(leucine codon number as indicated along X axis) | 15 (2), 46 (6), 68<br>(10), 141 (14),<br>201 (18) | Different combinations of these sites were chosen as indicated in the X-axis of each panel. |
| tRNA accommodation rate at CTA68 (CSAT model) | 0.0607s<sup>-1</sup> | Fit from experiment |
| tRNA accommodation rate at CTA141 (CSAT model) | 0.135s<sup>-1</sup> | Fit from experiment |
| tRNA accommodation rate at CTA201 (CSAT model) | 0.128s<sup>-1</sup> | Fit from experiment |
| tRNA accommodation rate at CTA15 (CSAT model) | 0.0658s<sup>-1</sup> | Fit from experiment |
| tRNA accommodation rate at CTA46 (CSAT model) | 0.0873s<sup>-1</sup> | Fit from experiment |
| tRNA accommodation rate at CTC68 (CSAT model) | 0.616s<sup>-1</sup> | Fit from experiment |
| tRNA accommodation rate at CTC141 (CSAT model) | 0.795s<sup>-1</sup> | Fit from experiment |
| tRNA accommodation rate at CTC201 (CSAT model) | 0.784s<sup>-1</sup> | Fit from experiment |
| tRNA accommodation rate at CTC15 (CSAT model) | 0.198s<sup>-1</sup> | Fit from experiment |
| tRNA accommodation rate at CTC46 (CSAT model) | 0.378s<sup>-1</sup> | Fit from experiment |
| tRNA accommodation rate at CTT68 (CSAT model) | 0.21s<sup>-1</sup> | Fit from experiment |
| tRNA accommodation rate at CTT141 (CSAT model) | 0.75s<sup>-1</sup> | Fit from experiment |
| tRNA accommodation rate at CTT201 (CSAT model) | 0.566s<sup>-1</sup> | Fit from experiment |
| tRNA accommodation rate at CTT15 (CSAT model) | 0.177s<sup>-1</sup> | Fit from experiment |
| tRNA accommodation rate at CTT46 (CSAT model) | 0.289s<sup>-1</sup> | Fit from experiment |
| tRNA accommodation rate at CTA68 (SAT model) | 0.186s<sup>-1</sup> | Fit from experiment |
| tRNA accommodation rate at CTA141 (SAT model) | 0.479s<sup>-1</sup> | Fit from experiment |
| tRNA accommodation rate at CTA201 (SAT model) | 0.435s<sup>-1</sup> | Fit from experiment |
| tRNA accommodation rate at CTA15 (SAT model) | 0.214s<sup>-1</sup> | Fit from experiment |
| tRNA accommodation rate at CTA46 (SAT model) | 0.296s<sup>-1</sup> | Fit from experiment |
| tRNA accommodation rate at CTC68 (SAT model) | 3.0s<sup>-1</sup> | Fit from experiment |
| tRNA accommodation rate at CTC141 (SAT model) | 4.52s<sup>-1</sup> | Fit from experiment |
| tRNA accommodation rate at CTC201 (SAT model) | 4.63s<sup>-1</sup> | Fit from experiment |
| tRNA accommodation rate at CTC15 (SAT model) | 0.73s<sup>-1</sup> | Fit from experiment |
| tRNA accommodation rate at CTC46 (SAT model) | 1.56s<sup>-1</sup> | Fit from experiment |
| tRNA accommodation rate at CTT68 (SAT model) | 0.779s<sup>-1</sup> | Fit from experiment |
| tRNA accommodation rate at CTT141 (SAT model) | 3.78s<sup>-1</sup> | Fit from experiment |
| tRNA accommodation rate at CTT201 (SAT model) | 3.05s<sup>-1</sup> | Fit from experiment |
| tRNA accommodation rate at CTT15 (SAT model) | 0.66s<sup>-1</sup> | Fit from experiment |
| tRNA accommodation rate at CTT46 (SAT model) | 1.1s<sup>-1</sup> | Fit from experiment |
| tRNA accommodation rate at CTA68 (TJ model) | 0.0461s<sup>-1</sup> | Fit from experiment |
| tRNA accommodation rate at CTA141 (TJ model) | 0.0975s<sup>-1</sup> | Fit from experiment |
| tRNA accommodation rate at CTA201 (TJ model) | 0.0893s<sup>-1</sup> | Fit from experiment |
| tRNA accommodation rate at CTA15 (TJ model) | 0.0524s<sup>-1</sup> | Fit from experiment |
| tRNA accommodation rate at CTA46 (TJ model) | 0.066s<sup>-1</sup> | Fit from experiment |
| tRNA accommodation rate at CTC68 (TJ model) | 0.264s<sup>-1</sup> | Fit from experiment |
| tRNA accommodation rate at CTC141 (TJ model) | 0.289s<sup>-1</sup> | Fit from experiment |
| tRNA accommodation rate at CTC201 (TJ model) | 0.281s<sup>-1</sup> | Fit from experiment |
| tRNA accommodation rate at CTC15 (TJ model) | 0.151s<sup>-1</sup> | Fit from experiment |
| tRNA accommodation rate at CTC46 (TJ model) | 0.206s<sup>-1</sup> | Fit from experiment |
| tRNA accommodation rate at CTT68 (TJ model) | 0.135s<sup>-1</sup> | Fit from experiment |
| tRNA accommodation rate at CTT141 (TJ model) | 0.271s<sup>-1</sup> | Fit from experiment |
| tRNA accommodation rate at CTT201 (TJ model) | 0.247s<sup>-1</sup> | Fit from experiment |
| tRNA accommodation rate at CTT15 (TJ model) | 0.129s<sup>-1</sup> | Fit from experiment |
| tRNA accommodation rate at CTT46 (TJ model) | 0.173s<sup>-1</sup> | Fit from experiment |

All other parameters have values shown in Table S6.

**Table S4:** Simulation parameters for Fig. 4C and Fig. S3.

| Parameter | Value | Note |
| --- | --- | --- |
| Stall site identity | CTC |  |
| Stall site locations as codon number along *yfp*<br>(leucine codon number as indicated along X axis) | 60 (8), 64 (9), 68<br>(10), 119 (11), 125<br>(12), 137 (13),<br>141 (14) | Different<br>combinations of<br>these sites were<br>chosen as<br>indicated in the X-axis of each figure. |
| tRNA accommodation rate at CTC68 (CSAT model) | 0.488s<sup>-1</sup> | Fit from experiment |
| tRNA accommodation rate at CTC119 (CSAT model) | 0.395s<sup>-1</sup> | Fit from experiment |
| tRNA accommodation rate at CTC125 (CSAT model) | 0.388s<sup>-1</sup> | Fit from experiment |
| tRNA accommodation rate at CTC137 (CSAT model) | 0.539s<sup>-1</sup> | Fit from experiment |
| tRNA accommodation rate at CTC141 (CSAT model) | 0.747s<sup>-1</sup> | Fit from experiment |
| tRNA accommodation rate at CTC60 (CSAT model) | 0.497s<sup>-1</sup> | Fit from experiment |
| tRNA accommodation rate at CTC64 (CSAT model) | 0.45s<sup>-1</sup> | Fit from experiment |
| tRNA accommodation rate at CTC68 (SAT model) | 2.22s<sup>-1</sup> | Fit from experiment |
| tRNA accommodation rate at CTC119 (SAT model) | 1.56s<sup>-1</sup> | Fit from experiment |
| tRNA accommodation rate at CTC125 (SAT model) | 1.61s<sup>-1</sup> | Fit from experiment |
| tRNA accommodation rate at CTC137 (SAT model) | 2.52s<sup>-1</sup> | Fit from experiment |
| tRNA accommodation rate at CTC141 (SAT model) | 4.13s<sup>-1</sup> | Fit from experiment |
| tRNA accommodation rate at CTC60 (SAT model) | 2.09s<sup>-1</sup> | Fit from experiment |
| tRNA accommodation rate at CTC64 (SAT model) | 1.8s<sup>-1</sup> | Fit from experiment |
| tRNA accommodation rate at CTC68 (TJ model) | 0.235s<sup>-1</sup> | Fit from experiment |
| tRNA accommodation rate at CTC119 (TJ model) | 0.199s<sup>-1</sup> | Fit from experiment |
| tRNA accommodation rate at CTC125 (TJ model) | 0.207s<sup>-1</sup> | Fit from experiment |
| tRNA accommodation rate at CTC137 (TJ model) | 0.239s<sup>-1</sup> | Fit from experiment |
| tRNA accommodation rate at CTC141 (TJ model) | 0.279s<sup>-1</sup> | Fit from experiment |
| tRNA accommodation rate at CTC60 (TJ model) | 0.243s<sup>-1</sup> | Fit from experiment |
| tRNA accommodation rate at CTC64 (TJ model) | 0.222s<sup>-1</sup> | Fit from experiment |

All other parameters have values shown in Table S6.

**Table S5:** Simulation parameters for Fig. 5.

| Parameter | Value | Note |
| --- | --- | --- |
| Stall site identity | CTA | Panel A, B |
| Stall site location (codon number along *yfp*) | 201 | Panel A, B |
| Threshold tRNA accommodation rate for selective abortive termination | 22s<sup>-1</sup> | Panel A, B |
| Threshold tRNA accommodation rate for non-selective abortive termination | 0s<sup>-1</sup> | Panel A, B |
| tRNA accommodation rate at stall site (TJ model) | 0.0893s<sup>-1</sup> | Panel A, B |
| tRNA accommodation rate at stall site (SAT model) | 0.435s<sup>-1</sup> | Panel A, B |
| tRNA accommodation rate at stall site (CSAT model) | 0.128s<sup>-1</sup> | Panel A, B |

All other parameters have values shown in Table S6.

**Table S6:** Parameters common to all simulations.

| Parameter | Value | Units | Reference |
| --- | --- | --- | --- |
| Cell volume | 10-18 | m<sup>-3</sup> | (54, 55) |
| Cell doubling time | 30 | min | This work |
| Concentration of tRNAs | 4.08 × 10<sup>5</sup> | molecules/cell | (54) |
| Rate of peptide-bond formation and translocation | ≈ 22 (1/0.045) | s<sup>-1</sup> | (54) |
| tRNA accommodation rate at normal sites (not stall sites) | 80–1650 | s<sup>-1</sup> | Inferred (Methods) |
| Ribosome footprint size on mRNAs | 10 | codons | This work |
| Number of distinct mRNA species | 2 | No units | This work |
| Ribosome-tRNA association rate constant | 2 × 10<sup>7</sup> | M<sup>-1</sup>s<sup>-1</sup> | (56, 57) |
| tRNA aminoacylation rate constant | 2 × 10<sup>10</sup> | M<sup>-1</sup>s<sup>-1</sup> | This work |
| Threshold tRNA accommodation rate for selective abortive termination | 22 | s<sup>-1</sup> | This work |
| Abortive termination rate constant (SAT and CSAT models) | 1 | s<sup>-1</sup> | This work |
| Abortive termination rate constant (TJ model) | 0 | s<sup>-1</sup> | This work |
| Translation initiation rate | 0.3 | s<sup>-1</sup> | This work |

**Table S7:**
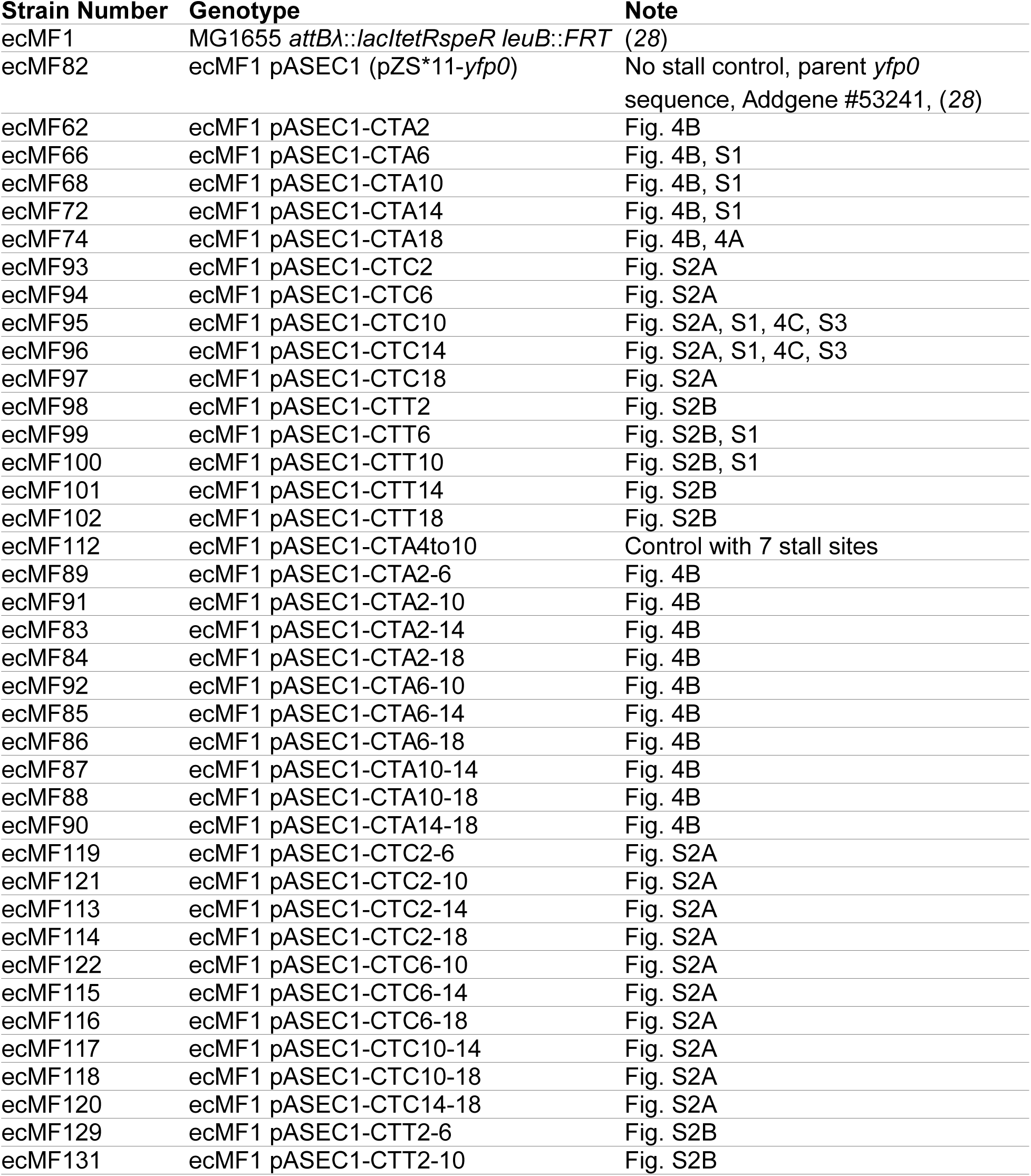
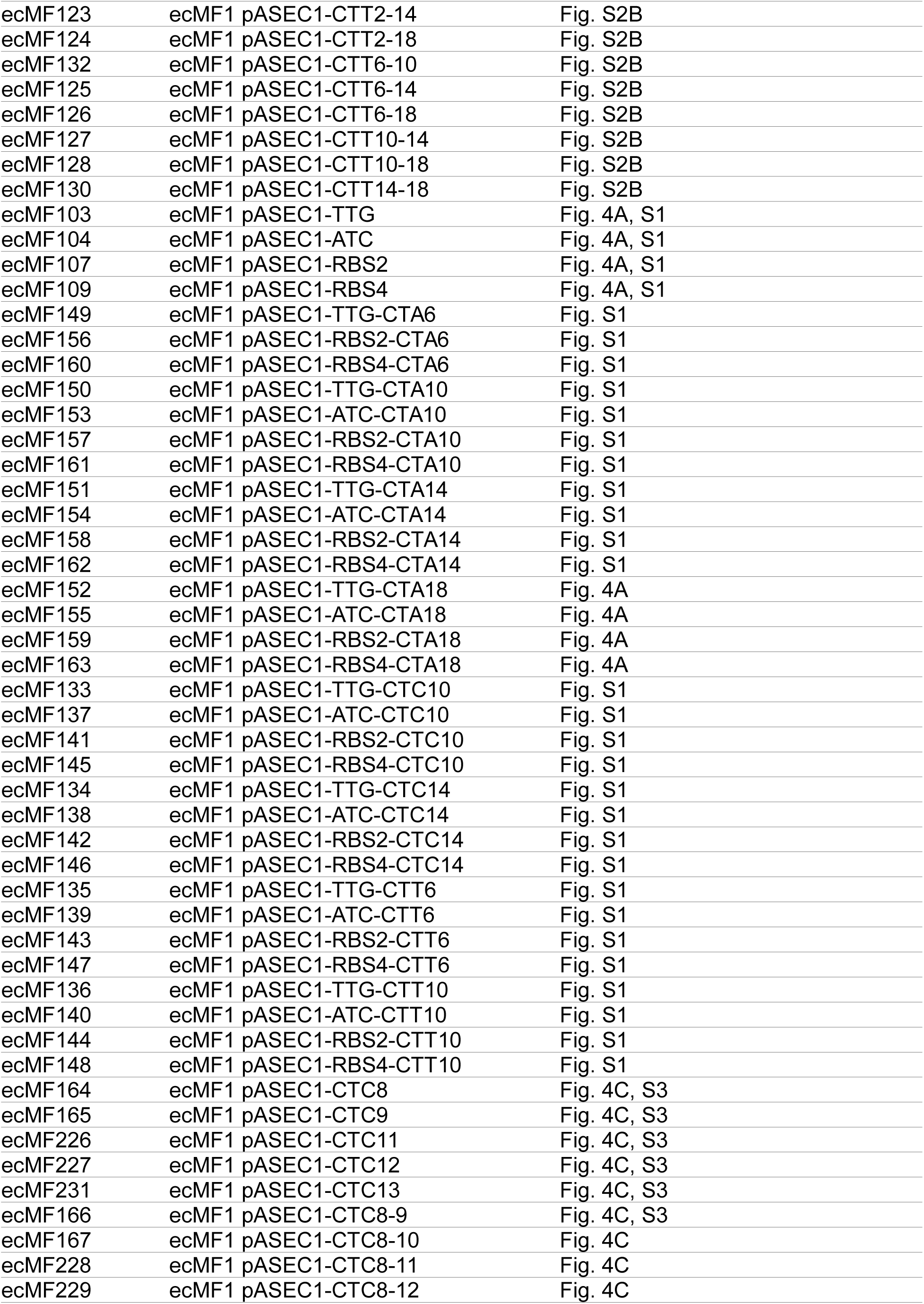
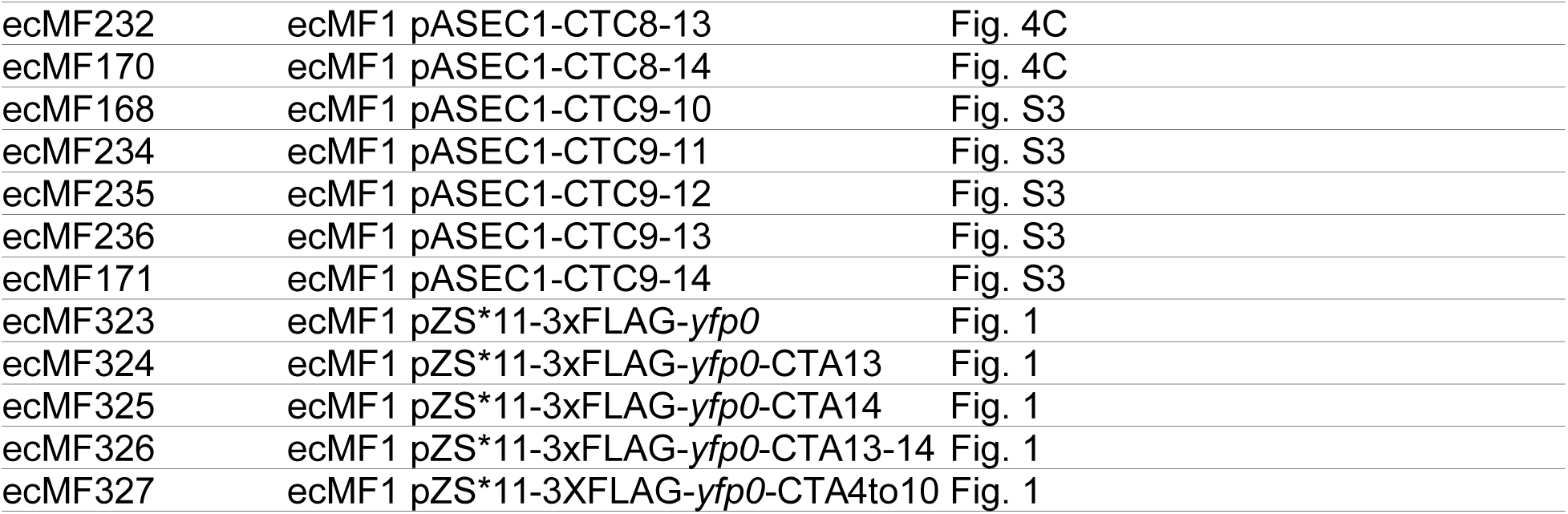
Strains and plasmids used in this study.

